# Characterizing co-expression networks underpinning maize stalk rot virulence in *Fusarium verticillioides* through computational subnetwork module analyses

**DOI:** 10.1101/237339

**Authors:** Man S. Kim, Huan Zhang, Huijuan Yan, Byung-Jun Yoon, Won Bo Shim

## Abstract

*Fusarium verticillioides* is recognized as an important stalk rot pathogen of maize worldwide, but our knowledge of genetic mechanisms underpinning this pathosystem is limited. Previously, we identified a striatin-like protein Fsr1 that plays an important role in stalk rot. To further characterize transcriptome networks downstream of Fsr1, we performed next-generation sequencing (NGS) to investigate relative read abundance and also to infer co-expression networks utilizing the preprocessed expression data through partial correlation. We used a probabilistic pathway activity inference strategy to identify functional subnetwork modules likely involved in virulence. Each subnetwork modules consisted of multiple correlated genes with coordinated expression patterns, but the collective activation levels were significantly different in *F. verticillioides* wild type versus the mutant. We also identified putative hub genes from predicted subnetworks for functional validation and network robustness studies through mutagenesis, virulence and qPCR studies. Our results suggest that these genes are important virulence genes that regulate the expression of closely correlated genes, demonstrating that these are important hubs of their respective subnetworks. Lastly, we used key *F. verticillioides* virulence genes to computationally predict a subnetwork of maize genes that potentially respond to fungal genes by applying cointegration-correlation-expression strategy.

## Introduction

Maize stalk rot is a complex disease, primarily caused by a series of fungal pathogens. Charcoal rot (by *Macrophomina phaseolina*), Fusarium stalk rot (by *Fusarium verticillioides*), Gibberella stalk rot (by *F. graminearum*) and Anthracnose stalk rot (by *Colletotrichum graminicola*) are the major stalk rots that devastate maize-growing regions in the US^1,2^. Losses due to stalk rot come in several different forms including stalk breakage, lodging, premature death of the plant, and the interruption of the normal grain filling process. Pathogens typically overwinter in the crop residue from the previous year and produce spores in the next growing season that will serve as the primary inoculum source. It is generally perceived that when crops experience abiotic stress, particularly at the end of the growing season, pathogens take advantage and colonize vulnerable stalk tissues^2-4^. But overall, we still lack a clear understanding of how these stalk rot fungi colonize and progress through pathogenesis.

To better understand the mechanism of pathogenesis, we screened for loss-of-virulence *F. verticillioides* mutants and identified a gene, *FSR1*, that is responsible for the deficiency^5^. Microscopic examination of inoculated stalks revealed the wild-type fungus vigorously colonizing vascular bundles and causing rot, whereas the mutant showed limited colonization and rot in stalks. *FSR1* encodes a protein that shares high similarity with striatins, a group of proteins found in eukaryotes that form complexes with kinases and phosphatases to regulate diverse cellular functions^6-8^. Recent studies have demonstrated important cellular and physiological roles of striatin proteins in *Sordaria macrospora, Neurospora crassa, Aspergillus nidulans*, and *C. graminicola*^9-12^. Our laboratory also revealed the importance of the coiled-coil motif of Fsr1 in virulence and demonstrated how Fsr1 forms a complex with other proteins to regulate stalk rot virulence^13,14^. These discoveries collectively support our hypothesis that Fsr1/striatin-mediated signal transduction plays a critical role in regulating stalk rot pathogenesis.

One of the intriguing questions we are aiming to answer is the impact of Fsr1 in cellular signaling associated with *F. verticillioides* virulence. To unravel the complex web of genetic interactions in *F. verticillioides* and maize, we decided to take advantage of next-generation sequencing (NGS) and explore the transcriptomic subtnetwork modules underpinning *FSR1*-mediated fungal virulence by computational network-based analysis. Our goal was to develop probabilistic and systematic models to investigate the interrelationship between genes rather than relying on quantitative comparison of transcript abundance as a measure of significance. Our NGS study was designed to capture dynamic changes in gene expression during maize stalk colonization by *F. verticillioides* wild type and *fsr1* mutant. To capture dynamic changes in transcriptome, samples were harvested from three distinct phases of stalk pathogenesis: establishment of fungal infection, colonization and movement into the vascular bundles, and host destruction and collapse^15^. A total of six independent biological replications were prepared and analyzed for each sample, since increasing the number of replicates was important for us to implement our computational analysis for identifying subnetwork modules that show strong differential expression.

As described in our previous work^15^, our strategy is to first construct the co-expression network of *F. verticillioides* using partial correlation, and search through these networks to detect subnetwork modules that are differentially expressed in the two *F. verticillioides* strains. Subsequently, we use the probabilistic pathway activity inference scheme^16^ to predict the activity level of potential subnetworks, followed by applying a computationally efficient branch-out technique to find the subnetworks that display the largest differential expression. Through this computational pipeline, we can identify potential pathogenic modules, which consist of genes that show coordinated behavior in *F. verticillioides* but also behaving differently in the wild type and the mutant. We can also screen for potential gene modules that contain orthologs of well-known virulence genes in other phytopathogenic fungi.

Biological functions, including virulence, are executed through elaborate collaboration of various biomolecules, and there has been increasing interest in the computational identification of functional modules from large-scale experimental data. In this study, we performed a comparative analysis of two distinct *F. verticillioides* RNA-Seq datasets, where one set was obtained from wild-type *F. verticillioides* and the other set from a loss-of-virulence *fsr1* mutant. For a systematic analysis of the infection transcriptome, we first predicted the co-expression network of the fungus. Subsequently, we identified functional subnetwork modules in the co-expression network consisting of interacting genes that display strongly coordinated behavior in the respective datasets. A probabilistic pathway activity inference method was adopted to identify three subnetwork modules likely to be involved in *F. verticillioides* virulence. Each subnetwork consisted of multiple genes with coordinated expression patterns, but more importantly we targeted subnetworks whose collective activation level is significantly different in the wild type versus the mutant. We then applied a series of mathematical criteria to predict the hub gene in each network and functionally tested their role in *F. verticillioides* virulence and the maintenance of network robustness.

## RESULTS

### NGS data preparation and relative expression analysis

We performed NGS using Illumina HiSeq 2000 and generated 36 independent libraries (*i.e*., six libraries per each time point - 3 dpi [infection], 6 dpi [colonization], and 9 dpi [rot] - for wild type and the *fsr1* mutant). For analysis and prediction in this study, we used 24 sample libraries from the last two time points (6 dpi and 9 dpi) to focus on gene regulation mechanism in the latter stages of maize-fungal interaction. Acquisition of read counts of all *F. verticillioides* genes was completed by mapping NGS reads to *F. verticillioides* strain 7600 reference genome^17^ using Bowtie2^18,19^ and Samtools^20^. Through filtering process, we eliminated genes with insignificant expression and therefore 9446 genes were selected for downstream analysis. We normalized the read counts of these genes by their corresponding gene length and also based on relative expression quantification against two *β*-tubulin genes (FVEG_04081 and FVEG_05512). Percentages of the two *β*-tubulin read abundance were traced over 24 replicates in 6 dpi & 9 dpi to examine their expression consistency. Mean (μ) and standard deviation (σ) of the percentages for β-tubulins were μ=0.058%, σ=0.0056 for FVEG_04081 and μ=0.035%, σ=0.0044 for FVEG_05512. The general information of our NGS datasets is shown in Fig. 1A. From these genes, we selected 324 most significantly differentially and highly expressed genes either in wild type or *fsr1* mutant from our datasets, where all replicates were normalized and analyzed for their individual relative expression levels at three different time points. As shown in a heat map with three distinct time points (Fig. 1D), 155 genes (red) are expressed significantly higher in the wild type and 169 genes (blue) are expressed significantly higher in *fsr1* mutant (Fig 1D. and Table S1). As explained earlier, the relative abundance was acquired by the two-step normalization by each gene length as well as *β*-tubulin genes, and was selected by *t*-test statistics score measurement (|*t*-test score| > 3.5). However, this common NGS analysis focuses on relative expression of individual genes but does not allow us to predict gene-gene associations and system-level changes across correlated genes during pathogenesis.

**Fig. 1.**
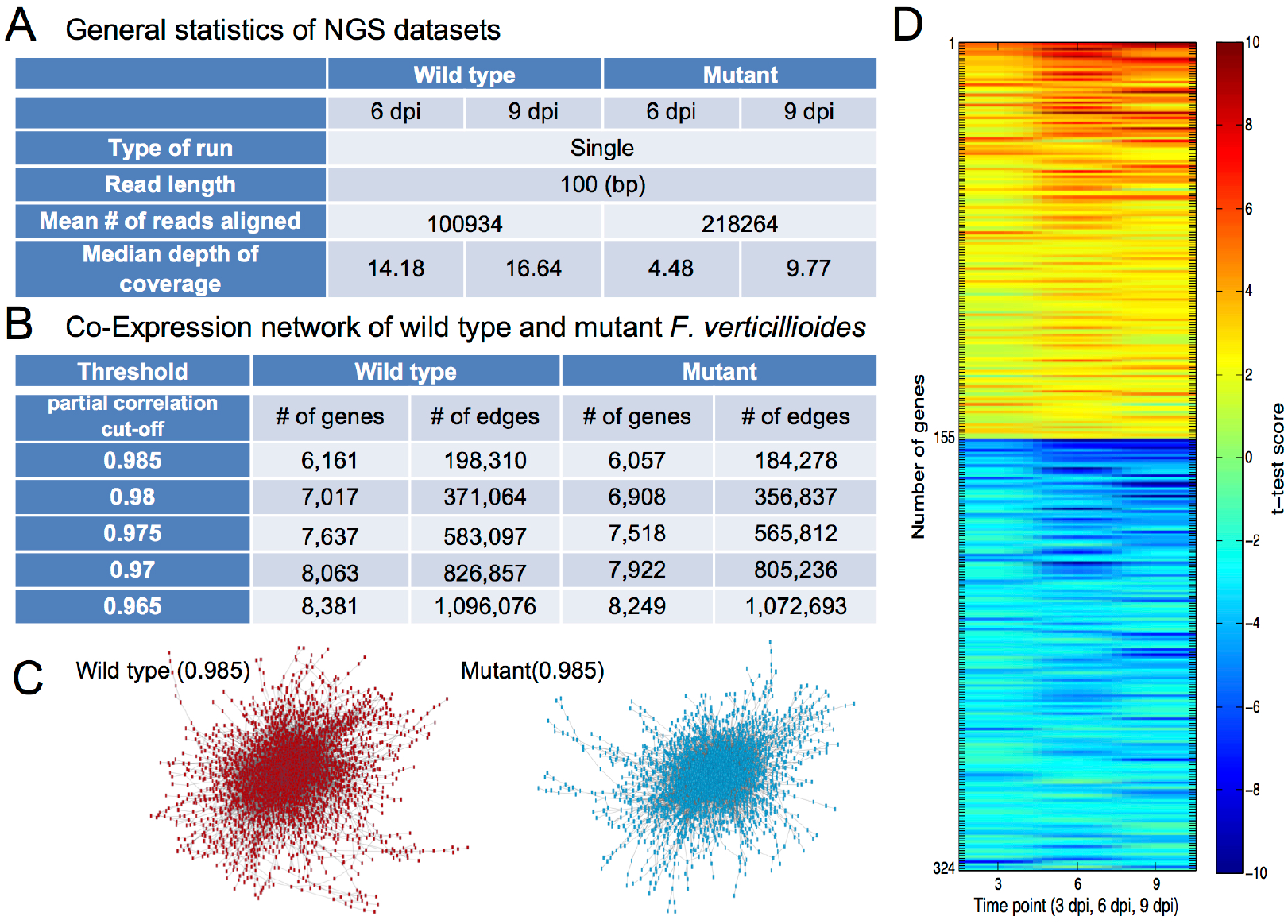
NGS statistics of significantly differentially expressed genes. A) General statistics of next-generation sequencing (NGS) datatsets. Reads from wild type and mutant (6 dpi and 9dpi) were mapped to *F. verticillioides* reference genome. B) Co-expression network of wild type and mutant *F. verticillioides*. We applied five distinct threshold levels to generate five different co-expression networks. The table shows number of genes and all possible edge combinations in each co-expression network. C) Schematic depiction of wild type and mutant co-expression networks at threshold level 0.985. D) The heat map provides a schematic overview of 324 most significantly differentially expressed genes at three distinct time points. A total of 155 genes are expressed significantly higher in the wild type (red) while a total of 169 genes are expressed significantly higher in the mutant (blue). In this selection, genes whose absolute *t*-test statistics score is higher than 3.5 were chosen, where the relative abundance was acquired by the two-step normalization which considers each gene length as well as relative expression against beta-tubulin genes over all the time points (3 dpi, 6 dpi, and 9 dpi). Next, *t*-test scores of the selected genes were again measured over relative expression levels normalized at each time point, and displayed in colors.

### Identification of *F. verticillioides* subnetwork modules

We developed a computational workflow that allows us to build co-expression networks from *F. verticillioides* NGS datasets^15^. We first inferred the co-expression networks for the wild type as well as the *fsr1* mutant utilizing the preprocessed gene expression data by using the partial correlation^21^ (Supplementary Information). In this co-expression network, we applied five distinct thresholds (0.965, 0.97, 0.975, 0.98, and 0.985), thereby constructing five different co-expression networks. The number of genes and edges between genes are shown in Fig. 1B. When these co-expression networks are illustrated with all member genes and possible edges, we can generate a complex web of scale-free networks (Fig. 1C). However, the aim of our proposed network-based NGS data analysis is to search through these co-expression networks to identify subnetwork modules that are differentially activated between the *F. verticillioides* wild type and *fsr1* mutant, that can considerably differ in terms of virulence potentials (Fig. 2).

**Fig. 2.**
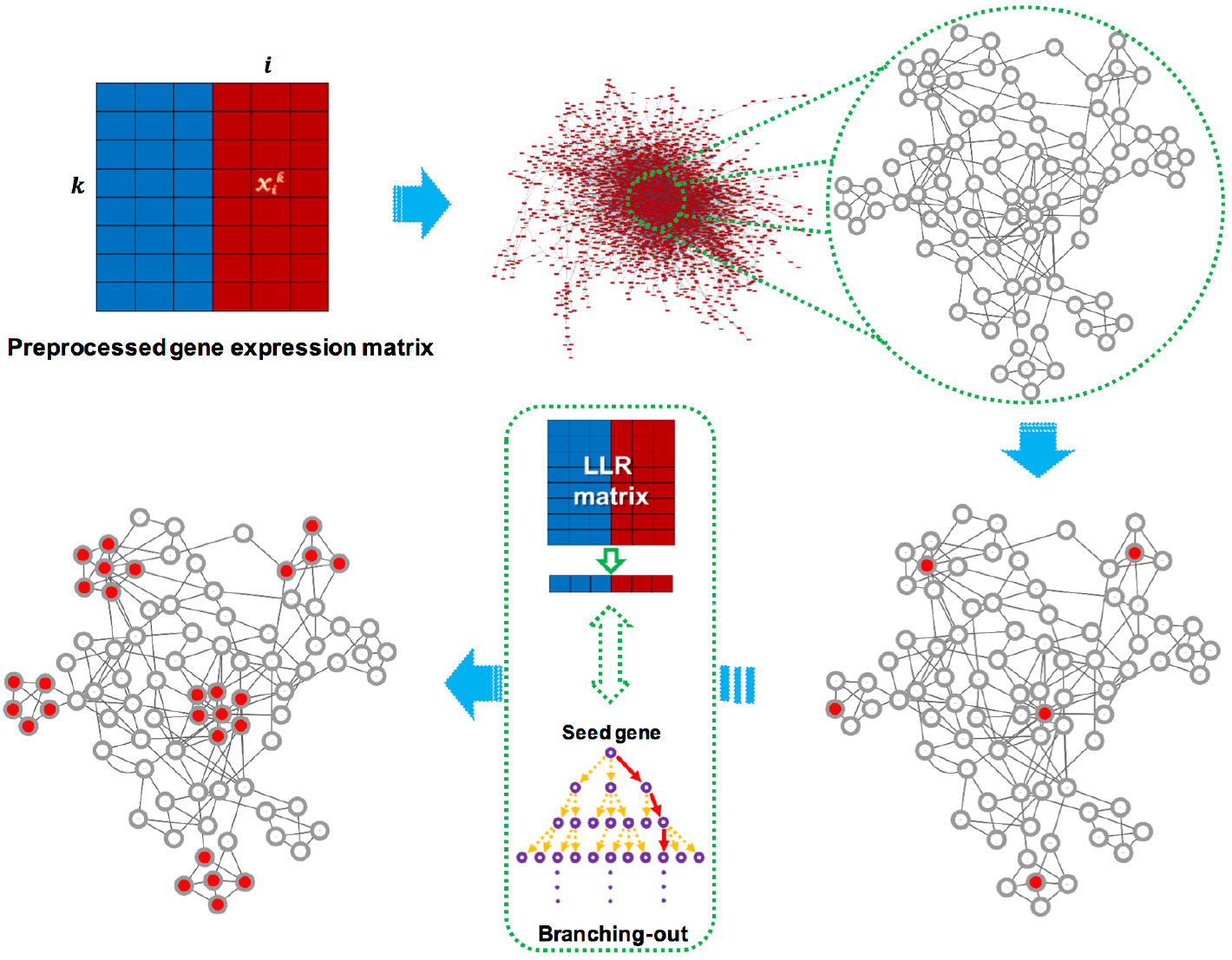
Schematic overview of our network-based NGS data analysis. Our aim is to search through large co-expression networks to identify subnetwork modules that are differentially activated between two different conditions (*e.g*. wild type versus mutant). Initial selection of seed genes (*i.e*. top 1% differentially expressed genes) is followed by a series of computational procedures described previously^15^. Through this process, we can identify subnetwork modules that show significant difference in virulence potentials.

By following this proposed strategy, we developed two potential subnetwork modules differentially activated during *F. verticillioides* pathogenesis. We performed additional analyses with six different characteristics for selecting hubs and their modules followed by our network-based comparative analysis approach^15^ (Fig. 3). In the subnetwork module fine-tuning process, two modules in Fig. 4 showed the minimum discriminative power increase for the entire module adjustment as 22% and 27% while over 90% of modules displayed smaller than 20% increase. Note that our approach probabilistically focuses on generating subnetwork modules whose member genes have high likelihood of showing associated expression patterns to each other across all replicates using the log-likelihood-ratio (LLR) matrix that demonstrates how likely each gene would express in *F. verticillioides* wild type or the mutant. As a result, our network-based computational analysis approach found potential subnetwork modules that show harmonious coordination of member genes as well as strong differential activity between the two strains.

**Fig. 3.**
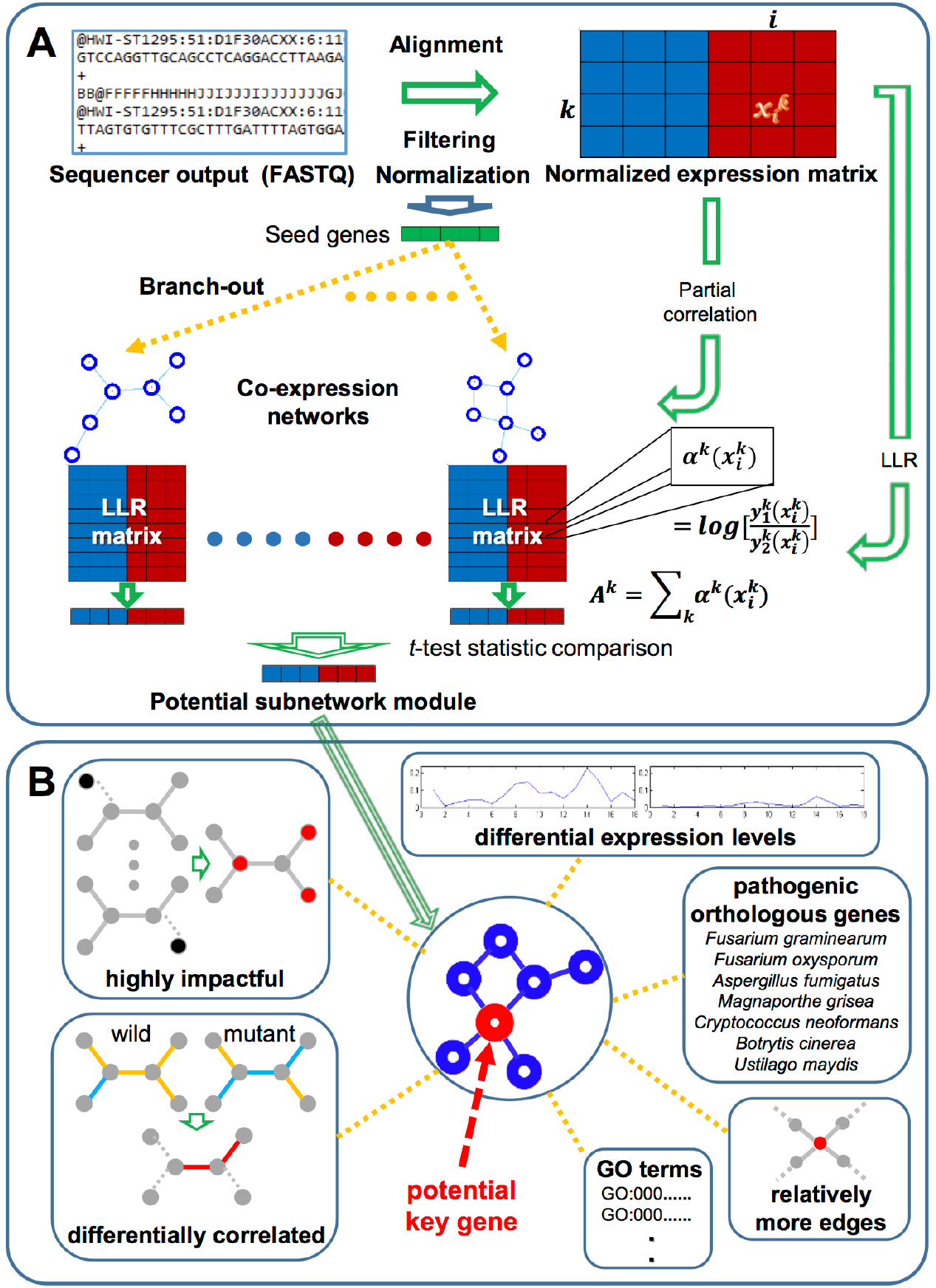
Computational prediction procedure for identifying key potential pathogenicity genes. A) Raw NGS datasets are preprocessed, *i.e*. alignment, filtering, and normalization, before they are applied for inferring *F. verticillioides* co-expression networks by means of partial correlation. In addition, these datasets were also converted into a log-likelihood ratio (LLR) matrix for downstream analysis. Next, subnetwork modules are extended from seed genes, significantly differentially expressed between the two different conditions (*F. verticillioides* wild type vs. mutant), as long as they keep sufficient strength of differential activity between the two strains [1-2]. B) Each potential hub (virulence-associated) gene is predicted in its detected subnetwork module based on several criteria: i) highly impactful in a probabilistic manner, ii) relatively differentially correlated between two strains (wild vs. mutant), iii) relatively more connected in the given module, iv) relatively significantly differentially expressed, v) orthologous to known pathogenicity-associated genes of other fungal species, vi) annotated to significant GO terms with other member genes. Through this proposed analysis approach, we identified potential functional genes showing significant differential activity between the two conditions as well as strong association with virulence.

**Fig. 4.**
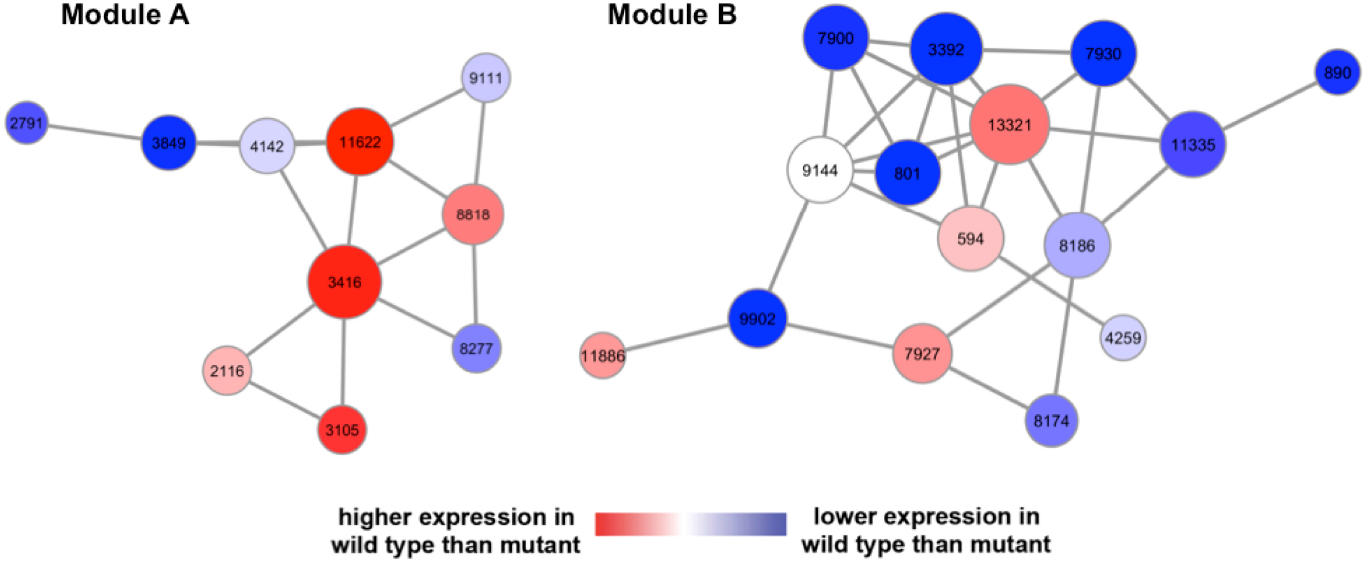
Potential subnetwork modules associated with the *F. verticillioides* pathogenicity. Our network-based comparative analysis identified three potential virulence-associated subnetwork modules differentially activated between the two strains (*F. verticillioides* wild type vs. mutant). Module A is composed of ten *F. verticillioides* genes, where 80% of them were annotated to a significant GO term GO:0044444 “cytoplasmic part”. Module B is comprised of fifteen genes, where four of them (FVEG_07930, FVEG_00890, FVEG_11886, and FVEG_00594) were annotated to a significant GO term GO:0006810 “transport”. The node sizes are directly proportional to their number of edges and the node colors vary according to their discriminative power measured by *t*-test scores.

### Computational characterization of two key *F. verticillioides* subnetwork modules

From our network-based comparative analysis, we identified two potential pathogenicity-associated subnetwork modules differentially activated between the wild-type and *fsr1* mutant strains (Fig. 4). Module A was composed of ten *F. verticillioides* genes, where 80% of these were annotated with a significant GO term cytoplasmic component (GO: 0044444) (http://biit.cs.ut.ee/gprofiler/index.cgi)^22^. However, it is important to note that majority of these genes have no known function and these GO functions were chosen solely based on predicted protein motifs (Table S2). Module B was comprised of fifteen genes, where four (FVEG_07930, FVEG_00890, FVEG_11886, and FVEG_00594) were annotated with a significant GO term transport (GO:0006810). The eleven other genes were hypothetical proteins with some knowledge of their functional domains. But this module showed relatively higher percentage of genes with no GO terms and no functional protein domains compared to module A (Table S2). Once we defined these subnetwork modules, we analyzed all member genes *in silico* to predict potential hub genes that may hold a key role in *F. verticillioides* pathogenicity.

In module A, we selected FVEG_11622 as a potential pathogenicity-associated hub gene based on following observations: i) FVEG_11622 deteriorated the differential probabilistic activity level of its given module from wild type to mutant by 26% (the mean of other member genes was 16%), ii) correlation coefficients of FVEG_11622 decreased from wild type to mutant by 0.26 and 0.34 for Pearson’s and Spearman rank, respectively (the mean of other member genes was 0.14 and 0.19), iii) FVEG_11622 contained four edges to other member genes (the mean of other member genes was 2.8), iv) FVEG_11622 demonstrated significant expression decrease from wild type to mutant (t-score of 4.4), and v) orthologous gene of FVEG_11622 in *Botrytis cinerea* (*BC1G*) is recognized as having a role in fungal virulence. The predicted hub gene FVEG_11622, which was tentatively designated as *FvEBP1*, encodes a putative 238-AA hypothetical protein that harbors Emopamil-binding protein (EBP) domain (pfam05241). In mammalian systems, this protein family is known to be associated with endoplasmic reticulum and plays a critical role in sterol isomerization and lipoprotein internalization^23^. An emopamil binding protein BcPIE3 in *Botrytis cinerea* which shares significant structural similarities to mammalian EBPs was shown to be important for virulence^24^. *FvEBP1* has four direct edges to FVEG_03416, FVEG_04142, FVEG_08818 and FVEG_09111. FVEG_03416 is an alginate lyase gene, and contains an alginate lyase domain which is important for fructose and mannose metabolism. FVEG_04142 is a V-type proton ATPase subunit F. V-type ATPases have hydrogen-exporting ATPase activity and are involved in ATP hydrolysis coupled proton transport. FVEG_09111 is a hypothetical protein, containing a PX-associated domain. The function of this protein is unknown, but its N-terminus is always found to be associated to a PX domain which is involved in targeting of proteins to cell membranes. FVEG_08818 is a hypothetical protein with a methyltransferase domain.

Using the same approach, we identified FVEG_00594 as the potential pathogenicity-associated hub gene in module B based on following observations: i) FVEG_00594 reduced the differential probabilistic activity level of its detected module from wild type to mutant by 24% (the mean of other member genes was 13%); ii) correlation coefficient difference of FVEG_00594 between wild type and mutant was 0.34 and 0.4 for Pearson’s and Spearman rank, respectively (the mean of other member genes was 0.24 and 0.2); iii) FVEG_00594 included four edges to other member genes (the mean of other member genes was 3.7); iv) the difference of expression level of FVEG_00594 was higher in wild type although it did not show high significance (*t*-score as 0.8), and v) the ortholog of FVEG_00594 in *F. graminearum* (FG) is recognized as having a role in fungal virulence. FVEG_00594, designated *FvSYN1*, encodes a putative 377 amino-acid protein that harbors two well-recognized domains: syntaxin N-terminal domain (cd00179) and SNARE domain (cd15849). In budding yeast, the SNARE protein complex is involved in membrane fusion and protein trafficking for new synthesis and recycling of organelles^25^. SNAREs were originally classified into v-SNAREs and t-SNAREs according to their vesicle or target membrane localization^26^. Syntaxins belong to t-SNARE proteins and are shown to play an important role in membrane fusion in eukaryotic cells^27,28^. Syntaxins are known as a family of membrane-associated receptors for intracellular transport vesicles. Syntaxin and SNAREs are also known to anchor these newly synthesized and recycled proteins to the cytoplasmic surface^29^. SNARE proteins play critical and conserved roles in intracellular membrane fusion in eukaryotic cells^30^. They were known to mediate membrane fusion during all trafficking steps of the intracellular communication process, including the secretory and endocytic pathways^31^. *FvSYN1* has four directly associated genes in the subnetwork module: FVEG_03392, FVEG_04259, FVEG_09144 and FVEG_13321. Three of these genes (FVEG_03392, FVEG_04259, FVEG_09144) encode hypothetical proteins with no known functional motif thus making it difficult to predict their role. While FVEG_13321 is a hypothetical protein, it does contain a fungal Zn_2_Cys_2_ binuclear cluster domain, which is typically found in the family of fungal zinc cluster transcription factors^32,33^.

### Functional characterization of predicted hub genes associated with virulence

To test our hypothesis that *FvEBP1* (FVEG_11622) and *FvSYN1* (FVEG_00594) are putative hub genes of subnetwork modules A and B, respectively, and that they are important for *F. verticillioides* virulence. We generated gene knockout mutants Δfvebp1 and Δfvsyn1 through homologous recombination following our standard split marker protocol^34^. Hygromycin B phosphotransferase (*HPH*) was used as the selective marker, and homologous recombination outcomes were confirmed by PCR (data not shown) and Southern blots (Fig. S1). We first compared vegetative growth of these mutants on synthetic media (PDA, V8 agar and defined medium agar). While Δfvsyn1 strain showed reduced colony growth, Δfvebp1 strain exhibited no growth defect (Fig. 5A). The mutant Δfvsyn1 showed restricted radial vegetative grow while exhibiting more dense and fluffier mycelial growth on solid media when compared to the wild type (Fig. 5B). When cultures were harvested from YEPD broth, we did not observe a significant difference in fungal mass production (Fig. 5C). For spore production on V8 plates, Δfvsyn1 produced significantly reduced spores when compared to other strains (Fig. 5D).

**Fig. 5.**
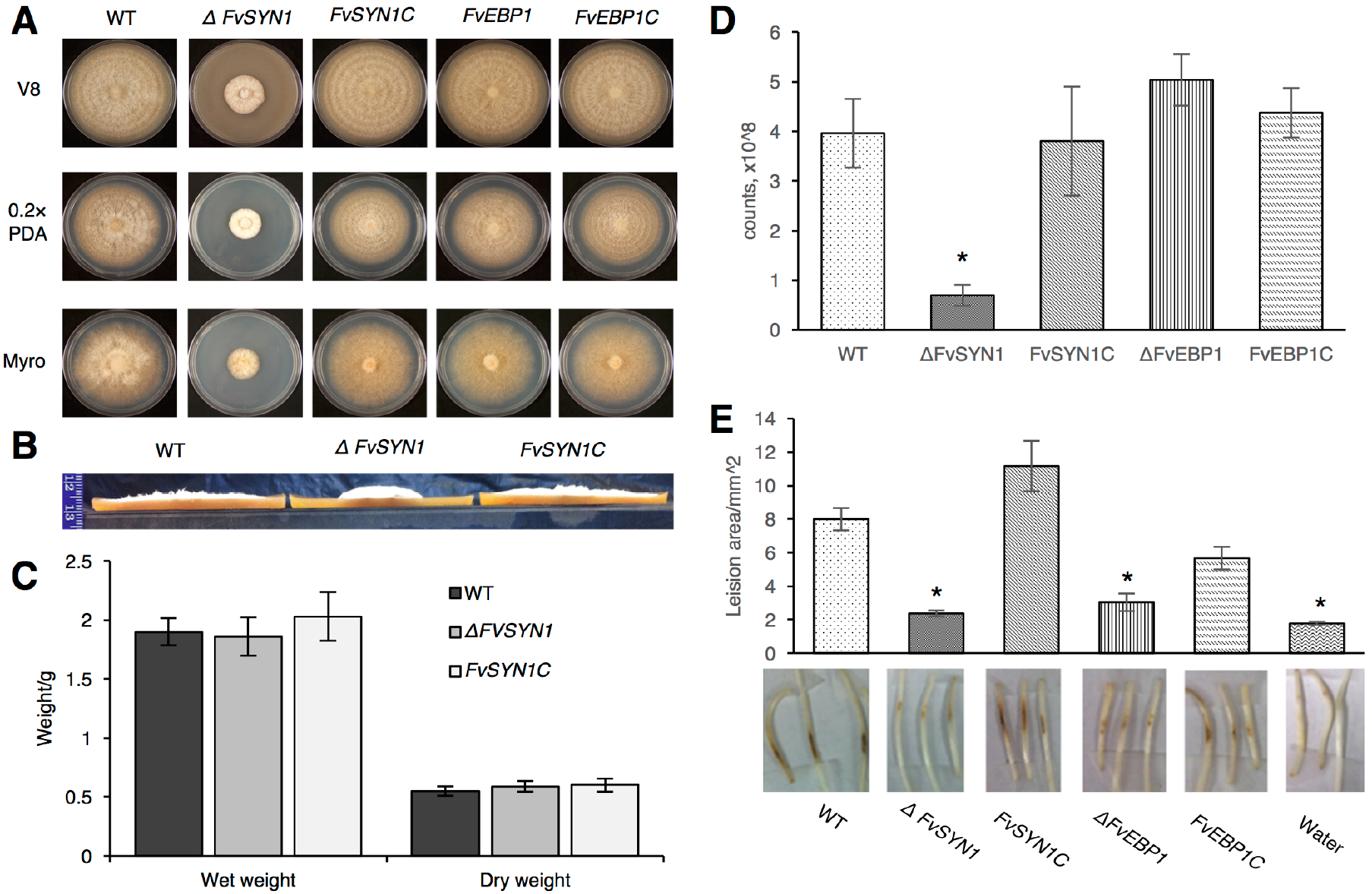
Functional characterization of *F. verticillioides FvSYN1* and *FvEBP1*. A) Vegetative growth of wild-type (WT), ΔFvsyn1, ΔFvebp1 and their complementation strains (FvSYN1C and FvEBP1C) were examined on V8, 0.2XPDA, and Myro agar plates. Strains were point inoculated with an agar block (0.5 cm in diameter) and incubated for 6 days at 25 °C under 14 h light/10 h dark cycle. B) Spores (2×10^7^) of WT, Δfvsyn1 and Fvsyn1C were inoculated in the center of V8 plates for 6 days at 25 °C under 14 h light/10 h dark cycle. Vegetative growth of WT, ΔFvsyn1 and FvSYN1C on V8 agar plates, strain growth condition was the same as described above. Agar plates were cut into half and pictures were taken from a side view. C) Fungal mass production of WT, ΔFvsyn1 and FvSYN1C strains was tested in YEPD broth. 100 μl spores (10^8^/ml concentration) were inoculated and incubated for 4 days at 25 °C and shaking at 150 rpm. Fungal mass production was quantified by weighing wet and dry fungal mass. The data presents the average and standard deviation of three independent experiments. D) Spore production of of WT, ΔFvsyn1, FvSYN1C, ΔFvebp1and FvEBP1C on V8 agar plates, strain growth condition was the same as described above. Spores were collected from agar plates and counted. E) One-week-old B73 seedlings were inoculated with 10^8^/ml spore suspension of fungal strains on mesocotyls. Lesion areas were quantified by Image J software after 2-week incubation. Asterisk above the column indicates statistically significant difference (P<0.05) analyzed by *t*-Test.

To test virulence, we inoculated B73 maize seedling mesocotyls with spore suspension of wild-type, Δfvebp1, and Δfvsyn1 strains (along with water as a negative control) following the previously described procedure^35^. When symptoms were observed after a 2-week incubation period, Δfvebp1 and Δfvsyn1 mutants showed significantly decreased levels of rot when compared with the wild-type progenitor (Fig. 4E). Mutants Δfvebp1 and Δfvsyn1 showed approximately 70% and 60% reduction in virulence when analyzed by average mesocotyl rot area (Fig. 5E). In order to test whether the mutant phenotype is due to a targeted gene replacement, we generated complementation strains of Δfvebp1 and Δfvsyn1 by co-transforming each mutant protoplasts with the respective wild-type gene (*FvEBP1* and *FvSYN1* with their native promoter and terminator) along with the geneticin-resistance gene. PCR was performed to confirm reintroduction of wild-type genes in complemented strains. FvSYN1C strain showed complete restoration of virulence on maize seedlings whereas FvEBP1C showed partial (~75%) recovery (Fig. 5E). These results suggested that *FvEBP1* and *FvSYN1* play an important role in virulence on maize seedling rot, and further convinced us that these two genes serve as the predicted hub gene of their respective subnetwork module.

### Testing network robustness in gene deletion mutants

A very important feature of these subnetwork modules is having robustness, *i.e*. the ability to respond to and withstand the external as well as internal stimuli while maintaining its normal behavior^36^. However, it is reasonable to predict that when we eliminate or disable a critical node (*i.e*. a hub gene), the network could be disrupted and shattered into isolated nodes. If a hub gene is eliminated from the subnetwork, we can hypothesize that other member genes, particularly those sharing direct edges, will exhibit disparate expression patterns.

We first tested correlated gene expression patterns in the wild type versus Δfvebp1 mutant by qPCR. We learned that gene expression levels of FVEG_03416, FVEG_04142, and FVEG_08818 were drastically lowered in the Δfvebp1 mutant than those observed in the wild-type progenitor (Fig. 6A). Furthermore, the FVEG_09111 gene expression level was not detectable in the mutant. Particularly, FVEG_04142 and FVEG_09111 showed higher levels of expression in the Δfsr1 mutant when compared to the wild type, and in Δfvebp1 that expression pattern is now reversed (Fig. 6B). These results show that when *FvEBP1* is no longer present in the subnetwork, expression levels of these genes, FVEG_03416, FVEG_04142, and FVEG_08818, and FVEG_09111, are drastically suppressed (Fig. 6A and B), suggesting *FvEBP1* is critical for proper regulation of these neighboring genes.

**Fig. 6.**
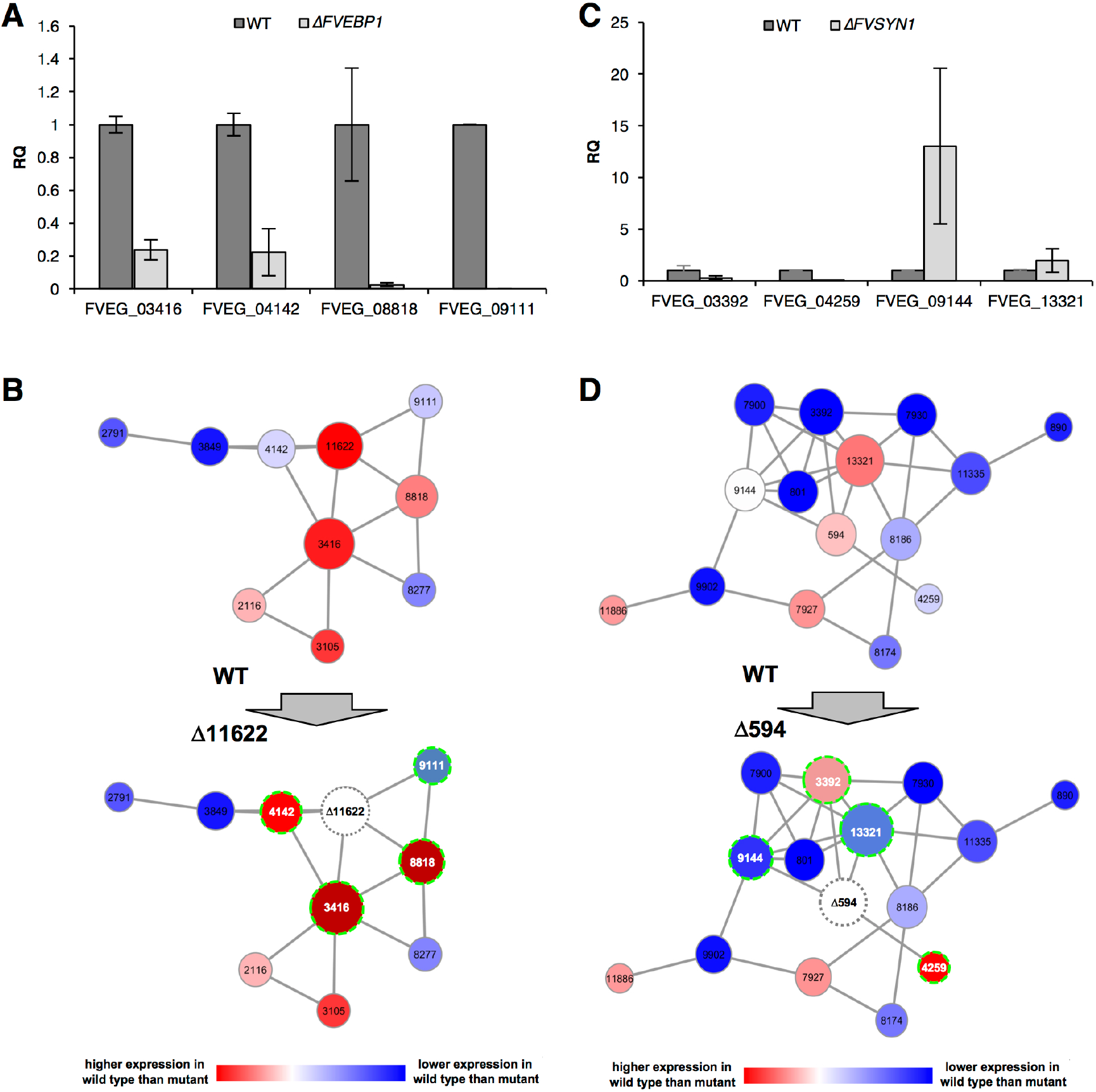
Altered expression of select neighboring genes as detected by qPCR. A) Relative quantification (RQ) of four neighboring genes (FVEG_3391, FVEG_9144, FVEG_13321, FVEG_4259) to predicted hub *FvSYN1* (FVEG_0594) in wild type (WT) versus ΔFvsyn1. RQ levels of four genes in WT were normalized to 1. B) Schematic overview of transcriptional changes of four neighboring genes (highlighted) of *FvSYN1* (FVEG_0594) observed in WT versus ΔFvsyn1. C) Relative quantification (RQ) of four neighboring genes (FVEG_4142, FVEG_8818, FVEG_3416, FVEG_9111) to predicted hub *FvEBP1* (FVEG_11622) in wild type (WT) versus ΔFvebp1. D) Schematic overview of transcriptional changes of four neighboring genes (highlighted) of *FvEBP1* (FVEG_11622) observed in WT versus ΔFvebp1.

In the Δfvsyn1 strain, we comparatively studied the expression pattern of four genes that directly share edges with *FvSYN1*. Three of the four genes tested, FVEG_03392, FVEG_04259 and FVEG_09144 showed a significant difference in expression levels between the wild type and Δfvsyn1 mutant. Significantly, FVEG_03392 and FVEG_04259, which showed lower expression level in the wild type when compared with Δfsr1 mutant, reversed its course and showed higher expression in the wild type when compared with Δfvsyn1 (Fig. 6C). FVEG_09144, which showed no difference in expression between wild type and Δfsr1, showed significantly higher expression in Δfvsyn1. FVEG_13321, which showed higher expression in wild type compared to Δfsr1, now exhibits statistically similar expression in wild-type and Δfvsyn1 (Fig. 6C and D). Collectively these data showed that *FvSYN1* and *FvEBP1* are important for regulating the expression of closely correlated genes, further providing evidence that these are important hub genes of their respective subnetworks.

## DISCUSSION

In this study, we assembled a streamlined computational network analysis pipeline to investigate the system-level coordinated changes across differentially activated genes rather than simply focusing on differential transcript abundance of individual genes, and to detect subtle processes that are not likely to be revealed by examining a small list of highly significant genes in this host-pathogen interaction. To generate meaningful prediction from limited datasets, comprehensive and rigorous investigation was needed. Thus, we mainly searched for comparable expression patterns probabilistically using a log-likelihood ratio matrix over replicates instead of just considering differential expression for identifying potential subnetwork modules. Also, we analytically investigated the given subnetwork modules with multidirectional analysis considering factors such as probabilistic impact, and differential correlation. Significantly, this comprehensive approach can help identify novel virulence-associated subnetwork modules as well as the key functional “hub” genes in fungal pathogens, such as *F. verticillioides*. This assembly of tool will be instrumental as we continue our effort to harness new and meaningful information from NGS data as we try to better understand complex pathosystems.

Our study mainly focused on analyzing the underlying transcriptional regulation in host-pathogen interactions. However, we do recognize that complex intercellular web of interactions in a living cell, not to mention between a host and its pathogen, are not limited to gene-gene association. Numerous constituents of the cell, *e.g*. DNA, RNA, protein, and metabolites, contribute to the structure and the dynamics of cellular network and ultimately behavior. However, in contrast to DNA and RNA, the resources available for us to generate systems-level proteome and metabolome datasets for network analyses are currently limited. In addition to this challenge, majority of host-pathogen systems have very limited genetic information available. For instance, as one can see from our three predicted modules majority of member genes encode hypothetical proteins with unknown, and vaguely predicted, functions. We primarily focused on developing this computational approach with the intent of investigating not-well defined biological systems with minimal bias toward existing genetic information, *i.e*. allocating higher scores toward known virulence genes in given species.

Furthermore, there is a greater challenge in refining subnetwork module development for host organisms that typically has larger and more complex genomes. Over 95% of our NGS data generated in this study came from maize, suggesting that maize stalk is actively responding to the pathogen invasion and colonization at transcriptional level^37^. In our earlier study, we developed a computational analysis pipeline, including the cointegration-correlation-expression approach, to predict potential maize defense-associated genes that show strong differential activation and coordination with known *F. verticillioides* virulence genes^37^. Here, we selected three *F. verticillioides* pathogenicity genes identified from our current work (FVEG_11622, FVEG_00594, and FVEG_09767 [FSR1]) to predict maize response subnetworks. An illustration of the potential subnetwork module associated with maize defense response against the *F. verticillioides* pathogenicity genes is shown in Fig. 7. We followed the procedure from narrowing down maize genes using the cointegration-correlation-expression approach to branching out potential defense-associated modules on maize co-expression networks^37^. The subnetwork module associated with maize defense system was composed of 28 maize genes, where genes relatively significantly expressed in wild type-infected are indicated in red and genes relatively significantly expressed in mutant type-infected are indicated in blue (Table S3). In this potential maize defense gene subnetwork module, we noticed that five maize genes (GRMZM2G102760_T01, GRMZM5G870932_T01, GRMZM2G001421_T02, GRMZM2G001696_T01, and GRMZM2G137535_T01) were annotated with a significant GO term defense response/incompatible interaction (GO:0009814), defined as “a response of a plant to a pathogenic agent that prevents the occurrence or spread of disease”. However, as seen in *F. verticillioides* subnetwork modules, we recognize that a large percentage of member genes encode hypothetical proteins and that transcriptional coordination does not always result in functional correlation. In addition, unlike *F. verticillioides* genes we characterized in this study, generating null mutants and performing network robustness assays are more strenuous for maize.

**Fig. 7.**
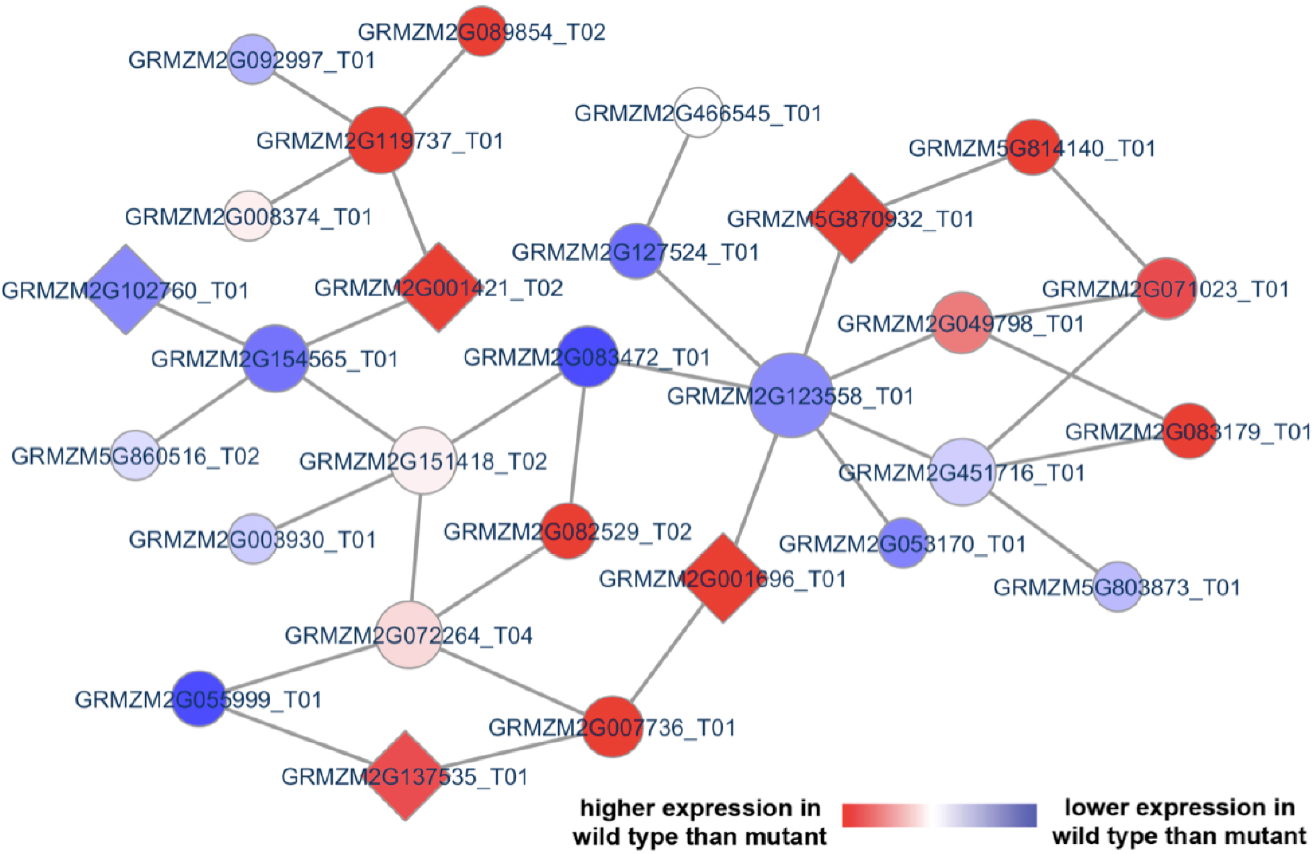
Potential subnetwork module associated with maize defense response. Our network-based comparative analysis of maize genes applied cointegration-correlation-expression strategy to identify a potential maize subnetwork module differentially activated between the two strains (*F. verticillioides* wild type vs. mutant). This predicted maize module is comprised of 28 member genes, and five were annotated to a significant GO term GO:0009814 “defense response, incompatible interaction”. These five genes were indicated by diamond-shaped illustration.

While difficulties mentioned above remain as obstacles, our effort demonstrates that the proposed network-based analysis pipeline can improve our understanding of the biological mechanisms that underlie host-pathogen interactions, and that it has the potential to unveil novel genetic subnetwork modules and hub genes critical for virulence in fungal pathogens. We are currently in the process of improving our computational pipeline with computational network querying that can estimate node correspondence probabilities to find novel functional pathways in biological networks^38-41^. This gene subnetwork approach can lead to the discovery of new quantifiable cellular subnetworks that can bridge the knowledge gaps in the maize-*F. verticillioides* system and be further applied to other plant-microbe pathosystems.

## METHODS

### Fungal strains, maize line, and RNA sample preparation

*F. verticillioides* strain 7600^17^ and fsr1 mutant^5^ were cultured at 25°C on V8 juice agar (200 ml of V8 juice, 3 g of CaCO3 and 20 g of agar powder per liter). Maize inbred B73, a progenitor of numerous commercial hybrids with no inherent resistance to stalk rot, was inoculated with *F. verticillioides* wild type and fsr1 mutant spore suspension as described previously^5,35^. Maize stalk samples were collected 3, 6, and 9 dpi using manual sectioning, and microscopically inspected to identify host tissue damage and/or fungal colonization, particularly in the vascular bundles. For each sample, sectioning was performed on at least three independent stalk samples from each stage of infection, and isolated tissues were pooled for RNA extraction with TRIzol reagent (Invitrogen). For each time point, we collected six pooled samples, thus thirty-six RNA samples in total. Standard QA/QC procedure for RNA samples was implemented at the Texas A&M AgriLife Research Genomics and Bioinformatics Service (College Station, TX) prior to sequencing.

### RNA Sequencing and data preprocessing

RNA sequencing was processed at the Texas A&M AgriLife Research Genomics and Bioinformatics Service using Illumina HiSeq 2000 as described previously^37^. We sequenced a total of 36 sample libraries, *i.e*. six libraries per each time point (3 dpi, 6 dpi, and 9 dpi) for wild type and the mutant inoculated maize stalks, but it is worth noting that, in this study, we only used sequencing data for the last two time points in this study (6 dpi and 9 dpi, hence 24 sample libraries in total) to focus on gene regulation mechanism in the latter stages of maize-Fusarium interaction. Next, acquiring read counts of all *F. verticillioides* genes was completed as described in our earlier reports^15,37^ by i) aligning the RNA-seq reads to the reference genome of *F. verticillioides* strain 7,600 obtained from the Broad Institute (http://www.broadinstitute.org) using Bowtie2^19^ and Samtools^20^, ii) filtering out genes insignificantly expressed over most of the replicates, thereby keeping 9446 genes for downstream analysis, iii) normalizing the read counts of these genes by their corresponding gene length and also based on expression levels of ***β***-tubulin genes (*i.e*. FVEG_04081 and FVEG_05512) to have relative expression quantification across all replicates.

### Prediction of *F. verticillioides* subnetwork modules associated with stalk rot

A procedure of identifying candidate functional subnetwork modules follows computational analysis pipeline described earlier^15^ with some modifications, and additional detail is provided in the Supplementary Information. From the preprocessed gene expression data, we performed conversion into log likelihood ratio (LLR) matrix, construction of co-expression networks tbased on partial correlation, and selection of the most significantly differentially expressed genes (*i.e*., top 1%) between the two strains (wild type vs. mutant) as seed genes. Based on this preparation, we applied computationally efficient branching out searching from a seed gene until it does not meet minimum discriminative power increase for each subnetwork module. The entire searching process was reiterated for every seed gene and for the five co-expression networks (Supplementary Information). Note that our approach probabilistically searches for subnetwork modules whose member genes have highly likely coordinated expression patterns to each other over all the replicates using the LLR matrix that demonstrates how likely each gene would be regulated in wild type or *fsr1* mutant. As a result, our network-based computational analysis approach found candidate subnetwork modules that show harmonious coordination of member genes as well as strong differential activity between the wild type and *fsr1* mutant.

### Prediction strategy for hub genes in each subnetwork module

Our proposed computational approach in this study simultaneously inferred potential *F. verticillioides* pathogenicity-associated hub or key functional genes in each subnetwork module while branching out the modules by performing multidirectional analysis on the detected candidate subnetwork modules with six different criteria, as depicted in Fig. 3B. First, we investigated how each gene is probabilistically impactful in its subnetwork module utilizing the probabilistic inference strategy applied in our previous work^15^. We estimated probabilistic differential activity of each gene by comparing discriminative power on the two phenotypes (wild type vs. *fsr1* mutant) between its given module with and without the gene. As described in our previous study^15^, we computed discriminative power difference (estimated by *t*-test statistics) between both activity levels (one with the gene and the other without the gene) by supposing ***ζ*** = {***g*_1_, *g*_2_,…, *g_n_***}, member genes in a subnetwork module, and ***e*** = (***e*^1^, *e*^2^,…, *e^n^***}, expression levels of the given genes. The discriminative power difference ***D***(***d***) was calculated as follows.

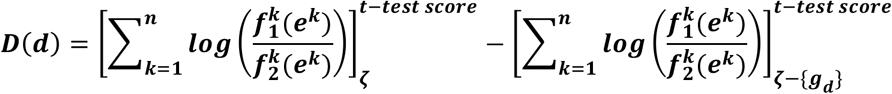

where 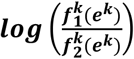 is the log-likelihood ratio (LLR), and ***f^k^***(***e***) is each conditional probability density functions (*i.e*., either wild or mutant). We subsequently considered genes whose discriminative power deterioration ***D***(***d***) relatively larger as candidate key genes. Second, we examined differential correlation of each gene with its connected gene in each module between the wild type versus the mutant using two correlation methods, *i.e*. Pearson’s correlation and Spearman rank correlation. We selected genes whose correlation coefficients with its neighboring genes were not only relatively significantly different between the two different networks, but also higher in the wild-type network as candidate hub genes. Third, we calculated number of edges of each gene to other member genes in each module since it is reasonable to predict that genes with more edges will exhibit more meaningful influence on the module. Fourth, we investigated expression difference of each gene in a given module between the two strains since we predict that altered expression of genes downstream of *FSR1* can follow that of *FSR1*. We selected genes that were significantly differentially expressed between the two conditions and relatively highly expression in in the wild type. Fifth, we listed orthologs of known pathogenicity genes in other well-studied fungal species such as *F. graminearum* (FG), *F. oxysporum* (FO), *Aspergillus fumigatus* (AF), *Botrytis cinerea* (BC1G), *Magnaporthe grisea* (MGG), *Ustilago maydis* (UM), and *Cryptococcus neoformans* (CNAG). We noted genes in a given module shown in the list of pathogenic genes as potential key genes. Finally, we considered whether each gene in a given module was associated with a significant GO term. We applied p-values of Benjamini-Hochberg false discovery rate (FDR) method^42^ to find the most relevant GO term to each gene and its given module based on g:Profiler (http://biit.cs.ut.ee/gprofiler/index.cgi)^22^. We chose genes annotated with a GO term that is the most significantly associated term with the given module as candidate key genes.

After identifying a candidate hub (or key functional) gene in its given subnetwork module, we performed additional fine-tuning procedure for more robust and reliable module prediction. Briefly, once we identified a candidate gene satisfying the abovementioned six criteria through subnetwork module extension process with 10% minimum discriminative power enhance, we performed module adjustment process by implementing the whole module extension for the given module as well as identifying the same potential gene by escalating the minimum discriminative power enhance by 1%. We repeatedly applied this fine-tuning process until the predicted hub (or key functional) gene did not meet all conditions. This process also stopped when the given subnetwork module grew smaller down to the arbitrary set minimum size of seven genes.

### Nucleic acid manipulation, polymerase chain reaction (PCR), and fungal transformation

Standard molecular manipulations, including PCR and Southern hybridization, were performed as described previously^34^. Fungal genomic DNA was extracted using the OminiPrep genomic DNA extraction kit (G Biosciences, Maryland heights, MO, USA). The constructs for transforming *F. verticillioides* were generated with a split-marker approach described earlier^34^. Briefly, DNA fragments of 5’ and 3’ flanking regions of each gene were PCR amplified from wild-type genomic DNA. Partial Hygromycin B phosphotransferase (*HPH*) gene (*HP*- and -*PH*) fragments were amplified from pBS15 plasmid. 5’ and 3’ flanking region fragments were then fused with PH- and -HP fragments by single-joint PCR, respectively. The single-joint PCR products were transformed into wild-type fungal protoplast. For complementation, respective wild-type genes driven by its native promoter was co-transformed with a geneticin-resistant gene (*GEN)* into mutant protoplasts. All primers used in this study were listed in Table S4. *F. verticillioides* protoplast were generated and transformed following standard protocol^34^ with minor modifications. Murinase (2 mg/ml) was replaced with Driselase (5 mg/ml) (Sigma, St Louis, MO, USA) in the protoplast digestion solution. Transformants were regenerated and selected on regeneration medium containing 100 μg/ml of hygromycin B (Calbiochem, La Jolla, CA, USA) and/or 150 μg/ml G418 sulfate (Cellgro, Manassas, VA, USA) as needed. Respective drug-resistant colonies were screened by PCR and further verified by Southern analysis.

### Maize infection assays

Maize seedling rot pathogenicity assay was performed on 2-week old maize inbred lines B73 seedlings as previously described^35^ with minor modifications. Briefly, 1x10^8^/ml spore suspensions in YEPD broth along with YEPD control were inoculated on maize B73 mesocotyls. Plant mesocotyls were first slightly wounded by a syringe needle about 3cm above the soil. A 5-μl spore suspension was applied to the wound site. The seedlings were immediately covered with a plastic cover to create a high moisture environment suitable for infection and colonization. The seedlings were collected and analyzed after a 2-week growth period in the dark room. At least three biological and three technical replicates were performed for each fungal strain.

### Expression analysis of subnetwork member genes linked to predicted hub genes

Total RNA extractions were conducted by using RNeasy plant mini kit (Qiagen) according to manufacturer’s specifications and was quantified by Nanodrop. RNA was converted into cDNA using the Verso cDNA synthesis kit (Thermo Fisher Scientific, Waltham, MA) following the manufacturer’s protocol. qRT-PCR analyses were performed using the SYBR Green Dynamo Color Flash qPCR kit (Thermo Fisher Scientific) on an Applied Biosystems 7500 Real-Time PCR system. The *F. verticillioides* β-tubulin gene (*TUB-2*) was used as the endogenous calibrator. The amplification data analysis was done according to the manufacturer’s protocol.

### Identification of potential maize defense subnetwork module

We followed our previous analysis strategy^37^ to identify potential subnetwork modules associated with maize defense response against *F. verticillioides* virulence genes. We began searching for maize modules possibly responsible for its defense mechanism through cointegration-correlation-expression analysis: i) Cointegration was applied to track an interrelationship of expression levels between maize and *F. verticillioides* over all replicates. We applied the Engle-Granger correlation method to measure single cointegrating relations and *p*-value <= 0.05 was used to identify candidate maize genes that appear to have significant association with the identified *F. verticillioides* pathogenicity-associated genes, ii) correlation was utilized to trace patterns of expression levels between maize and *F. verticillioides* over all replicates. We used Pearson’s correlation coefficients to estimate their linear relationship and condensed maize genes into candidates whose expression patterns are highly correlated with that of *F. verticillioides* pathogenicity-associated genes (*i.e*., *p*-value <= 0.005), iii) We considered expression levels of maize genes over replicates and filtered out insignificantly expressed maize genes (*i.e*., maize genes whose mean expression levels were in the bottom 20% or not expressed in at least one replicate). We adjusted *p*-values of the cointegration-correlation-expression approach to have 50% of candidate maize genes predicted from a given *F. verticillioides* pathogenicity-associated gene were also among the candidates inferred from other given *F. verticillioides* gene. Based on the maize genes narrowed down through the cointegration-correlation-expression analysis, we identified subnetwork modules associated with maize defense response using our network-based comparative analysis approach.

## Availability of Materials and Data

The datasets generated during and/or analyzed during the current study are available from the corresponding author on reasonable request.

## Author Contributions

Research concepts: M.K., H.Z., W.B.S., B.-J.Y.; experiments performed: M.K., H.Z., and H.Y.; manuscript editing: M.K., H.Z., W.B.S. All authors reviewed and approved the manuscript.

## Acknowledgments

This work was supported in part by the National Research Initiative Competitive Grants Program (No. 2007-35319-18334) and the Agriculture and Food Research Initiative (No. 2013-68004-20359) from the US Department of Agriculture. We would like to thank Mr. Trent LeMaster (Bioenvironmental Science, Texas A&M) and Mr. Jae-Young Jun (Electrical & Computer Engineering, Texas A&M) for excellent technical assistance.

